# Antigenicity, stability, and reproducibility of Zika reporter virus particles for long-term applications

**DOI:** 10.1101/2020.04.21.047241

**Authors:** J. Charles Whitbeck, Anu Thomas, Kathryn Kadash-Edmondson, Ariadna Grinyo i Escuer, Celine Cheng, Grant C. Liao, Frederick W. Holtsberg, M. Javad Aman, Graham Simmons, Edgar Davidson, Benjamin J. Doranz

**Affiliations:** Integral Molecular, Inc., 3711 Market St, Philadelphia, PA 19104; Vitalant Research Institute, 270 Masonic Ave, San Francisco, CA 94118; Integrated Biotherapeutics, 4 Research Court, Suite 300 Rockville, MD 20850

## Abstract

The development of vaccines against flaviviruses, including Zika virus (ZIKV) and dengue virus (DENV), continues to be a major challenge, hindered by the lack of efficient and reliable methods for screening neutralizing activity of sera or antibodies. To address this need, we previously developed a plasmid-based, replication-incompetent DENV reporter virus particle (RVP) production system as an efficient and safe alternative to the Plaque Reduction Neutralization Test (PRNT). As part of the response to the 2015-2016 ZIKV outbreak, we recently developed pseudo-infectious ZIKV RVPs by modifying our DENV RVP system. The use of ZIKV RVPs as critical reagents in human clinical trials requires their further validation using stability and reproducibility metrics for large-scale applications. In the current study, we validated ZIKV RVPs using infectivity, neutralization, and enhancement assays with monoclonal antibodies (MAbs) and human ZIKV-positive patient serum. ZIKV RVPs are antigenically identical to live virus based on binding ELISA and neutralization results and are nonreplicating based on the results of plaque formation assays. We demonstrate reproducible neutralization titer data (NT_50_ values) across different RVP production lots, volumes, time frames, and laboratories. We also show RVP stability across experimentally relevant time intervals and temperatures. Our results demonstrate that ZIKV RVPs provide a safe, rapid, and reproducible reagent for large-scale, long-term studies of neutralizing antibodies and sera, which can facilitate large-scale screening and epidemiological studies to help expedite ZIKV vaccine development.

**AUTHOR SUMMARY:** ZIKV is a mosquito-borne virus that can cause severe birth defects and other disorders. Large outbreaks of ZIKV occurred in 2015 and 2016 and there are still no drugs or vaccines available to protect against ZIKV infection. Vaccine development has been hindered by the lack of safe and efficient methods to screen ZIKV neutralizing properties of patient sera or antibodies, especially in the context of large clinical trials. To address this unmet need, we developed and validated the use of ZIKV reporter virus particles (RVPs), a safe, high-throughput and quantitative alternative to using live virus for neutralization studies. We show that ZIKV RVPs are stable, show lot-to-lot consistency and provide reproducible neutralization data that is suitable for large-scale studies are needed for development of a ZIKV vaccine, epidemiologic surveillance, and high-throughput screening.

## INTRODUCTION

Zika virus (ZIKV) is an emerging tropical arbovirus that was first identified in humans in 1952 in Uganda and the United Republic of Tanzania (1). ZIKV outbreaks are possible wherever *Aedes* mosquito species are present, but ZIKV can also be transmitted sexually or through blood transfusion (2). Although infected individuals are often asymptomatic, ZIKV infection in adults can cause Guillain–Barré syndrome, a potentially fatal autoimmune disease characterized by muscle weakness and paralysis. Infection during pregnancy can result in congenital Zika syndrome in newborns, characterized by microcephaly, eye defects, deafness, and growth deficits (3, 4). The most recent ZIKV epidemic in 2015-2016 involved up to 1.3 million cases in Brazil alone, with disease rapidly spreading throughout South and Central America and to the Caribbean Islands (5-7).

ZIKV is an enveloped flavivirus with a single-stranded, positive-sense 10.6 kDa RNA genome that encodes seven non-structural and three structural proteins: capsid, premembrane (prM), and envelope (E) (8). E and prM are the immunodominant proteins for flaviviruses, including ZIKV and dengue virus (DENV). Identified anti-ZIKV antibodies predominantly target the E protein, although some important MAbs target non-structural protein 1 (NS1). Although the overall structure of the ZIKV E protein is similar to that of DENV and other flaviviruses, numerous ZIKV-specific structural features contribute to its distinct antigenicity (9-14).

The development of flavivirus vaccines continues to be a challenge hindered by the lack of efficient and reliable methods for screening human sera for functional antibodies. Historically, the Plaque Reduction Neutralization Test (PRNT) was the standard measure of flavivirus infectivity (15). The PRNT assay determines viral neutralization based on the decrease in the formation of viral plaques on a cell monolayer. However, PRNT has numerous disadvantages – it is relatively slow, uses a large amount of serum or antibody, can be highly variable, uses live infectious virus, and some strains of virus do not readily form plaques (16-19). To overcome many of these limitations, we and colleagues previously developed a plasmid-based, replication-incompetent DENV reporter virus particle (RVP) production system for DENV studies (20-22). Antigenically identical to wild-type viruses, RVPs incorporate virus-specific capsid and prM/E proteins, contain a modified RNA genome, and express a reporter gene upon cellular infection, providing an efficient, reproducible, and safe alternative to plaque assays.

Recently, as part of the international response to the ZIKV outbreak, we developed and optimized pseudo-infectious ZIKV RVPs by modifying our DENV RVP system (20). Our ZIKV RVPs have already been used for measuring endpoints in preclinical studies of two different ZIKV vaccines and for supplementing clinical testing of human serum samples (23-25). However, the use of ZIKV RVPs as critical reagents in human clinical trials requires their further validation using stability and reproducibility metrics for large-scale applications. Here, we validate our ZIKV RVPs using infectivity and neutralization assays with monoclonal antibodies (MAbs) and human ZIKV-positive serum, comparing data for reproducibility within experiments, across days, between RVP production lots, and across different laboratories. Stability was tested up to 37°C and after multiple freeze-thaw cycles. Finally, we compared RVP neutralization titers with those obtained from PRNT. Our results demonstrate that ZIKV RVPs provide a safe, rapid, and reproducible reagent for large-scale screening applications, which can facilitate screening and epidemiological studies and help expedite ZIKV vaccine development.

## MATERIALS AND METHODS

### Plasmids, Cells Lines, and Reporter Virus Particles

ZIKV RVPs were produced in BHK cells using the DENV RVP system described previously (20-22) except that the CprME structural genes for ZIKV (strain SPH2015) were substituted instead of DENV CprME. For all experiments, frozen ZIKV RVPs (stored at −80°C) were thawed for 3 minutes in a 37°C water bath and then placed on ice before use. The following cell lines were used in this study: K562, BHK, HEK-293T, Vero (all from ATCC), Raji-DC-SIGN-R (kindly provided by Robert Doms), and BHK cells expressing DC-SIGN (BHK21 clone 15, Center for Vector borne Diseases, UC Davis). All cell lines were grown at 37°C at 5% CO2 in DMEM complete medium (DMEM with 10% FBS and 1% penicillin-streptomycin, 2 mM L-glutamine, 10 mM HEPES, 1 mM sodium pyruvate and 1% MEM non-essential amino acids) (Corning). Raji-DC-SIGN-R cells were grown in RPMI with 10% FBS and 1% penicillin-streptomycin and 2 mM L-glutamine.

### Monoclonal Antibodies and Human Sera

The following antibodies were used in ZIKV binding and neutralization studies: DENV neutralizing pan-flavivirus MAbs 4G2 (ATCC), 1N5, and 4E8 (26), DENV and ZIKV neutralizing MAbs 1C19 (26), C8 (27), and C10 (27), and ZIKV neutralizing MAbs LM-081 (Integral Molecular), A9E (28), ZIKV-117, ZIKV-195, and ZIKV-116 (29). MAbs that do not bind ZIKV, CHIKV-specific MAb CKV063 (30) and DENV1-specific 1F4 (31), were used as negative controls. ZIKV positive IgG serum was obtained from Boca Biolistics (donor D000013779). Human sera were heat inactivated for 30 minutes at 56°C) prior to use. A polyclonal anti-prM antibody (GeneTex), and anti-E MAb D11C (D004) (32) were used in western analyses of RVPs.

### ZIKV RVP Infectivity Assays

Luciferase ZIKV RVPs were serially diluted (two-fold) using infection medium and dispensed into a 96-well black plate coated with Poly-D-Lysine so that the first well contained 50 µl of undiluted RVPs. “Infection Media” consists of complete DMEM supplemented with 10% FBS adjusted to pH 8 ± 0.05 with NaOH, used within 14 days of preparation. 3×10^4^ cells (BHK-DC-SIGN and Vero cells) or 4×10^4^ cells (Raji-DC-SIGN-R cells) were added to each well and plates were incubated for 72 h at 37°C. Following incubation, plates were spun for 10 minutes at 2,000 rpm (581g, Sorvall Legend XTR), medium was removed, and cells were lysed using 25 µl of freshly prepared 1X *Renilla* luciferase assay lysis buffer. Covered plates were incubated at room temperature with shaking at 500 rpm for 30 minutes. Luciferase activity (relative luminescence units, RLU) was assessed by adding 30 µl luciferase assay buffer (buffer + substrate) per well and gently mixing by manually swirling the plate (a 5 min incubation prior to reading the plate is recommended) before reading luciferase activity on an Envision plate reader (Perkin Elmer). Due to time sensitivity of the luciferase signal, plates were read in the same order as substrate addition. Luciferase assay reagents tested included the Promega *Renilla*-Glo Luciferase Assay System (E2710) (recommended), Promega *Renilla* Luciferase Assay System (E2810), Abcam Luciferase Reporter Assay Substrate Kit (ab228546), and Pierce *Renilla* Luciferase Glow Assay Kit (ThermoFisher 1616). To determine the Z’ value of the RVP infectivity assay, we infected BHK-DC-SIGN cells on one plate with 12 replicate serial dilutions (7-fold) of luciferase ZIKV RVPs from a single lot (P-258B). As a negative control, cells were incubated without ZIKV RVPs.

### ZIKV RVP Neutralization Assays

MAbs or sera were diluted in 250 µl Infection Media in a 96 well V bottom plate. MAb stocks were initially diluted to 120 µg/ml and 3-fold dilutions were made thereafter using aerosol barrier tips. Serum samples were initially diluted 5-fold and then 3-fold dilutions were made thereafter. 90 µl of diluted MAbs were transferred in duplicate to a black, 96-well plate pre-coated with Poly-D-Lysine. ZIKV RVPs were thawed and diluted to a working concentration using Infection Media. 90 µl of diluted RVPs were added to each well of the neutralization plate containing MAbs. The RVP volume added to each well was determined from the infectivity assay to provide a signal of 200,000 to 500,000 RLU per well on an Envision plate reader. Neutralization plates were covered and placed in a 37°C tissue culture incubator. Following a 1 hour incubation, 3×10^4^ BHK-DC-SIGN cells were added to each well in 50 ul of infection medium. Neutralization plates were incubated at for 72 hours at 37°C and luciferase activity was assessed as described above.

### ZIKV Plaque Reduction Neutralization Test (PRNT)

Endpoint 50% PRNT titers were performed essentially as previously described (33). MAbs or serum were diluted as described above for RVP assays and then incubated for 1 hour at 37°C with approximately 80 PFU of ZIKV (SPH2015). Vero cells were seeded the previous day in 6-well tissue culture plates at a density of 6 × 10^5^ cells/well. Medium was aspirated and replaced with the antibody/virus mixture, and plaques were allowed to develop under agar overlays. After 72 hours, 2 mL of 0.6% agar containing 50 ug/mL neutral red was overlaid and plaques were counted after an additional 72 hours.

### Plaque Formation Assay to Detect Replicating Virions in RVPs

Vero cells were seeded in 6-well tissue culture plates (3.5×10^5^ cells per well). After 1 day of growth, medium in wells was aspirated and replaced with ZIKV RVPs (1 mL/well) or live DENV2 virus diluted in BHK Infection Medium. Virus or RVPs were allowed to infect or attach to cells for 1 h at 37°C. An additional 1 mL of medium was added to each well, and cells were grown for 4 d at 37°C, 5% CO_2_. Cell monolayers were fixed in ice-cold methanol, blocked with 10% NGS in PBS +/+, and stained for Flavivirus Env glycoprotein using MAb 4G2 (2 ug/mL) followed by goat anti-mouse Alexa Fluor 488.

### Antibody-Dependent Enhancement (ADE) of Infection

ZIKV RVPs were mixed with serial dilutions of MAb IgG in black PDL-coated 96-well plates and incubated for 1 h at 37°C. K562 target cells were added to each well and incubated for 72 h at 37°C. Culture medium was removed, cells were lysed, and plates were analyzed for luciferase activity.

### Shotgun Mutagenesis Epitope Mapping

Epitope mapping was performed by shotgun mutagenesis essentially as described previously (34). A ZIKV Env expression construct (strain ZikaSPH2015) was subjected to high-throughput alanine scanning mutagenesis to generate a comprehensive mutation library (29). Each residue within prM/E was changed to alanine, with alanine codons mutated to serine. In total, 672 ZIKV prM/E mutants were generated (100% coverage), sequence confirmed, and arrayed into 384-well plates. Each Env mutant was transfected into HEK-293T cells and allowed to express for 22 hrs. Cells were fixed in 4% (vol/vol) paraformaldehyde (Electron Microscopy Sciences), and permeabilized with 0.1% (wt/vol) saponin (Sigma-Aldrich) in PBS plus calcium and magnesium (PBS++). Cells were incubated with purified MAbs diluted in PBS++, 10% normal goat serum (NGS) (Sigma), and 0.1% saponin. Primary antibody screening concentrations were determined using an independent immunofluorescence titration curve against wild-type ZIKV prM/E to ensure that signals were within the linear range of detection. Antibodies were detected using 3.75 µg/ml AlexaFluor488-conjugated secondary antibody (Jackson ImmunoResearch Laboratories) in 10% NGS/0.1% saponin. Cells were washed 3 times with PBS++/0.1% saponin, followed by 2 washes in PBS. Mean cellular fluorescence was detected using a high throughput flow cytometer (HTFC, Intellicyt). Antibodies with quaternary epitopes were screened in a similar way but in cells co-expressing prME and furin and without fixation. Antibody reactivities against each mutant Env clone were calculated relative to wild-type Env protein reactivity by subtracting the signal from mock-transfected controls and normalizing to the signal from wild-type Env-transfected controls. Mutations within critical clones were identified as critical to the MAb epitope if they did not support reactivity of the test MAb, but supported reactivity of other ZIKV antibodies. This counter-screen strategy facilitates the exclusion of Env mutants that are locally misfolded or have an expression defect (35).

### MAb Binding

To demonstrate that MAb LM-081 is specific for ZIKV, expression constructs for the wild-type prM/E envelope proteins of DENV1 (WestPac), DENV2 (16803), DENV3 (CH53489), DENV4 (TVP360), ZIKV (SPH2015), and West Nile virus (WNV NY99) were transfected into HEK-293T cells and allowed to express for 22 hrs. Binding by MAb LM-081 was then tested by flow cytometry, as described above.

### ELISA for ZIKV RVPs Using Human MAbs

To test RVPs by ELISA, 384-well white plates were coated with capture MAb 4G2 overnight, followed by washing with PBS and blocking with 3% BSA. RVPs or RVP production medium was incubated for 2 hours. Primary antibodies were incubated for 1 hour at a concentration of 1, 2, or 4 ug/ml, followed by addition of rabbit anti-human HRP secondary antibody diluted to 1:5000 (Southern Biotech). Primary and secondary antibodies were incubated in the presence of 3% BSA blocking buffer. Chemiluminescent reagent (Pierce Femto reagent for ELISA) was added and the plate was read on an Envision plate reader.

## RESULTS

### ZIKV RVPs are infectious and antigenically identical to live ZIKV

Flavivirus RVPs are replication-incompetent viral particles that contain a subgenomic reporter replicon (encoding luciferase or GFP) packaged by flavivirus capsid (C), premembrane/membrane (prM/M), and envelope (E) proteins (**Fig. 1A**). RVP infection of permissive cells can be quantitatively measured by luminescence or flow cytometry, depending on the reporter. We produced ZIKV RVPs using the DENV RVP system as previously described (20-22), but substituting the CprM/E genes from ZIKV strain SPH2015. The RVPs used here were produced with a *Renilla* luciferase reporter gene to enable high sensitivity detection of infection and facilitate rapid, microplate-based detection.

**Figure 1.**
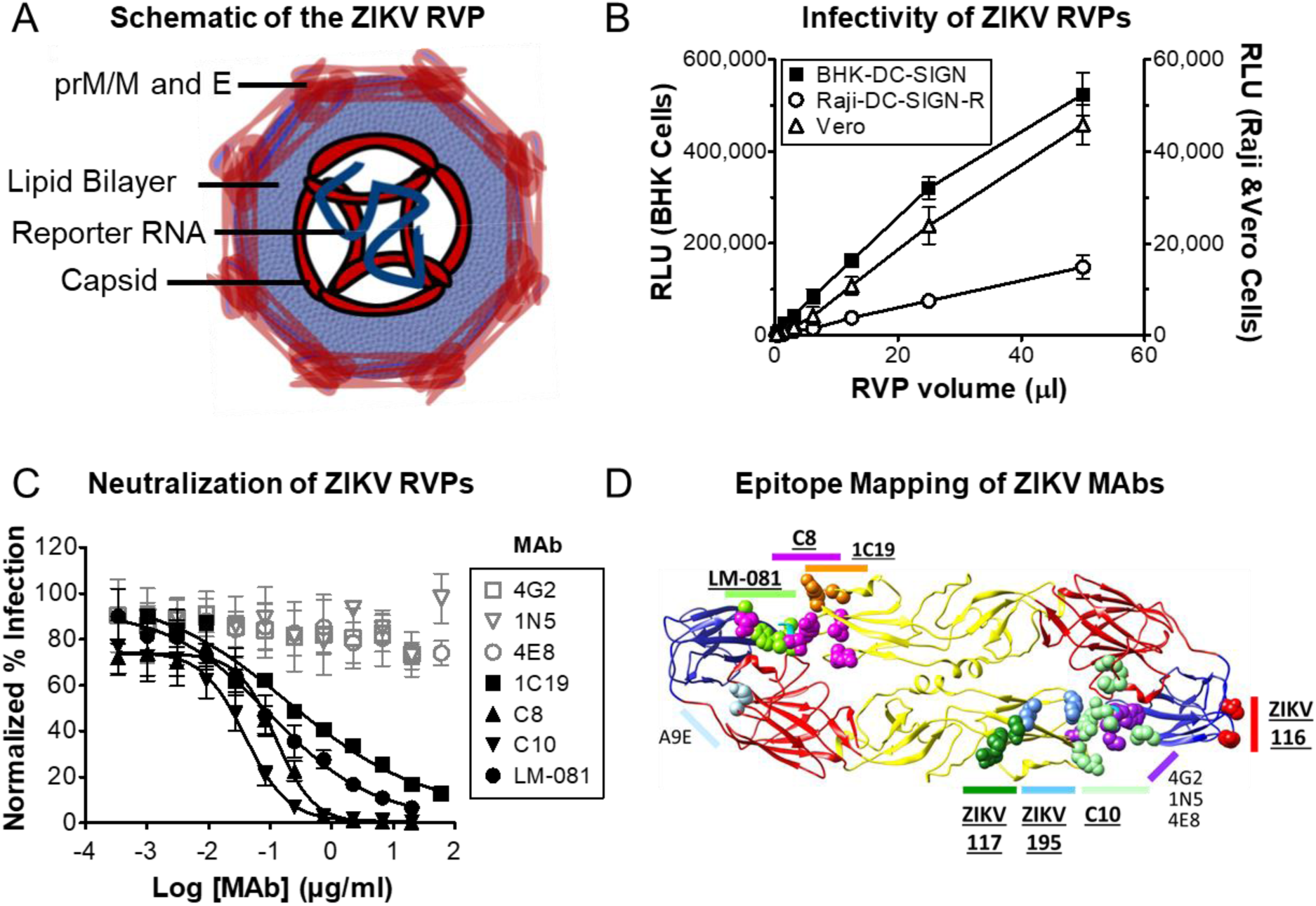
ZIKV RVPs are infectious and antigenically equivalent to live Zika virus. **(A)** Schematic of the ZIKV RVP, composed of capsid, prM/M, and E proteins from a defined ZIKV strain (SPH2015), and an RNA reporter genome replicon. **(B)** Luciferase ZIKV RVPs were tested for infectivity with cell lines that are commonly used for flavivirus infectivity studies, including BHK-DC-SIGN, Vero, and Raji-DC-SIGN-R cells. At 72 h after infection, cells were lysed and analyzed for luciferase activity (RLU). Values for BHK-DC-SIGN cells (filled symbols) are plotted on the left y-axis. Values for Raji-DC-SIGN-R and Vero cells (open symbols) are plotted on the right y-axis. All values represent n = 3 replicate wells, and error bars represent SD. Regression analysis indicates a linear relationship for the three datasets with RVP volume (R^2^ > 0.99). **(C)** Serial dilutions of the indicated MAbs were incubated with ZIKV RVPs for 1 h, followed by infection of BHK-DC-SIGN cells. After 72 h, cells were lysed and analyzed for relative luminescence units (RLU). All neutralization results are shown as normalized % infection (% of luciferase signal in the absence of MAb). For 1C19, 1N5, and 4E8, n = 2 and error bars indicate range. For all other data sets n = 7 or 8 and error bars represent SD. Gray symbols represent non-neutralizing MAbs, black symbols represent neutralizing MAbs. **(D)** Epitopes of 11 anti-ZIKV MAbs used in this paper, mapped onto the ZIKV structure. MAbs that neutralize ZIKV are indicated in bold and underlined.

To test the infectivity of ZIKV RVPs, Baby hamster kidney (BHK) cells expressing DC-SIGN, which acts as an attachment factor for DENV and ZIKV (36, 37), were infected with ZIKV RVPs. We obtained a linear relationship between the luciferase signal and the volume of RVPs added (**Fig. 1B**). We also tested four different luciferase assay detection systems to determine which was best suited to measuring RVP infectivity (**Fig. S1**). The Promega *Renilla*-Glo Luciferase Assay System (catalog #E2810) generated the most stable signal, with minimal change over 50 min after an initial 5 min incubation, so we recommend this system for high-throughput RVP neutralization experiments. ZIKV RVPs expressing luciferase resulted in a Z-factor (Z’) (38) score of 0.79, indicating an excellent signal separation for high-throughput screening, with low variability between infected and non-infected wells.

We also tested the infectivity of luciferase ZIKV RVPs in two other cell lines: Vero cells, and Raji cells expressing DC-SIGN-R (Raji-DC-SIGNR). Vero cells are commonly used for ZIKV studies and were recently used in conjunction with our ZIKV RVPs to determine serum neutralizing antibody titers in preclinical studies of a candidate ZIKV vaccine (23). We previously used Raji-DC-SIGNR cells to validate our DENV RVPs (20). As expected, all cell lines were infected by the ZIKV RVPs (**Fig. 1B**). BHK-DC-SIGN cells supported the highest levels of luciferase activity, so were chosen for subsequent experiments.

ZIKV RVPs are directly formed by the CprME structural proteins of ZIKV so are designed to be antigenically identical to native virus. As an initial test of the antigenicity of ZIKV RVPs, we assayed antigen-specific neutralization of ZIKV RVPs (**Fig. 1C**) using several MAbs with distinct conformational epitopes (**Fig. 1D**). 1C19 is a pan-flavivirus MAb that neutralizes ZIKV and DENV by binding residues in the b-c loop near the fusion loop (26). C8 and C10 are highly conformational, cross-reactive MAbs that neutralize ZIKV and DENV by binding to a quaternary “envelope dimer epitope” (EDE) region (39). LM-081 is a ZIKV-specific (**Fig. 2B**) potent neutralizing MAb generated and sold by Integral Molecular that binds a quaternary epitope across Domain II (DII) and Domain III (DIII) of adjacent ZIKV E protein monomers. MAbs 4G2, 1N5, and 4E8 are pan-flavivirus MAbs that neutralize DENV by binding epitopes in the fusion loop (26) but are known not to neutralize ZIKV (40, 41). As predicted, ZIKV RVPs were neutralized by 1C19, C8, C10, and LM-081, but not by control MAbs 4G2, 1N5, and 4E8.

**Figure 2.**
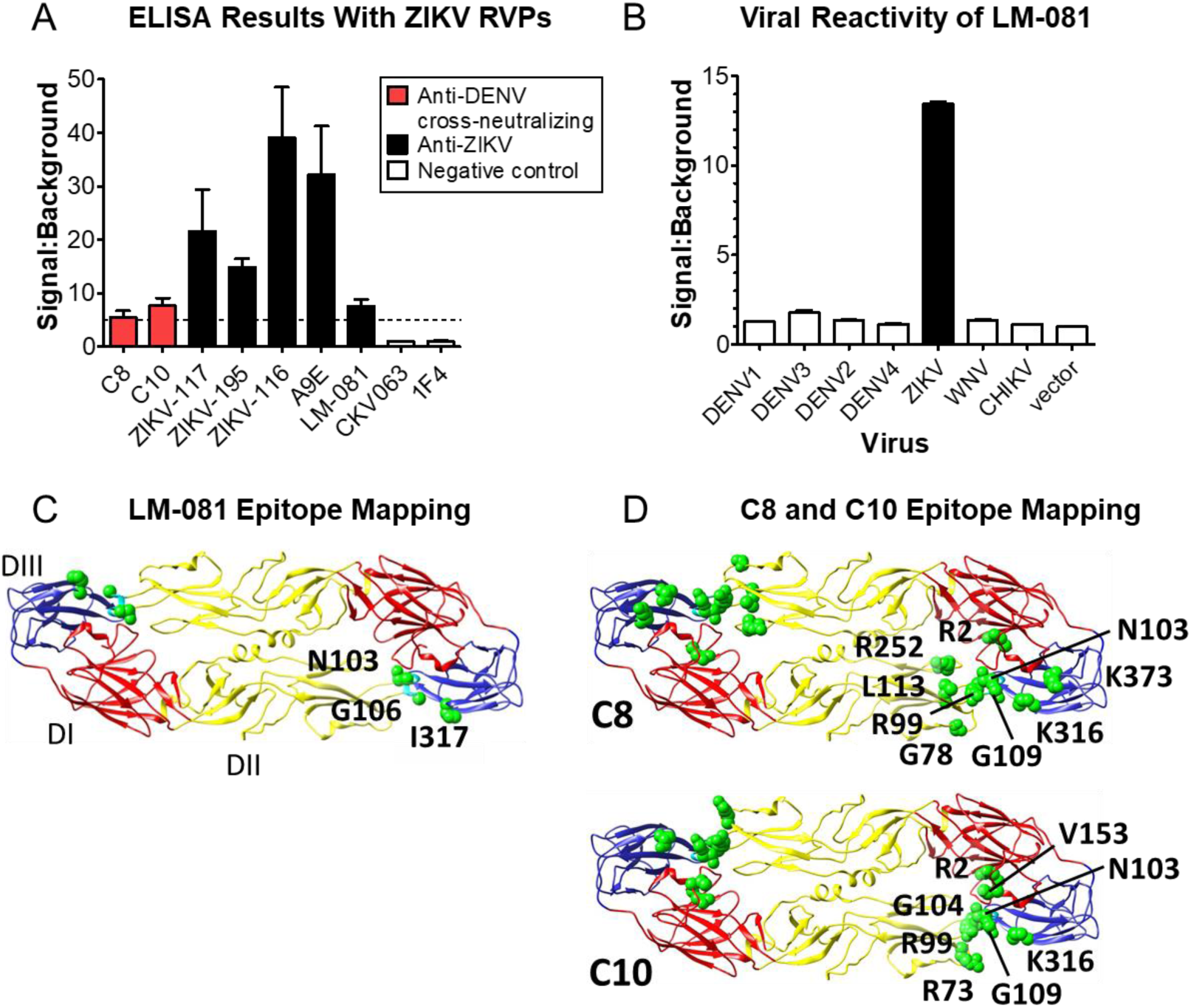
ZIKV RVPs are antigenically equivalent to live virus. **(A)** ELISA was performed using ZIKV RVPs with various conformational MAbs. Detection MAbs were two anti-DENV MAbs with ZIKV cross-neutralizing activity (C8, C10), five neutralizing anti-ZIKV MAbs (ZIKV-117, ZIKV195, ZIKV116, A9E, LM-081), and two negative control MAbs (CHIKV-specific CKV063 and DENV1-specific 1F4). The capture MAb was a mouse anti-fusion loop MAb (4G2). Mean signal to background ratio (S:B) of three concentrations of detection MAb (1, 2, and 4 μg/mL) is shown with the SD (error bars). Dashed horizontal line represents S:B = 5. (**B)** Flow cytometry analysis of LM-081 binding to viral envelope proteins expressed in human cells shows that LM-081 binds specifically to ZIKV and not to other flaviviruses, including DENV serotypes 1-4. **(C)** Shotgun mutagenesis epitope mapping of LM-081 revealed 2 critical residues in the fusion loop on one E monomer (N103, G106), and one critical residue on Domain III of an adjacent monomer (I317). **(D)** Shotgun mutagenesis epitope mapping of anti-DENV EDE MAbs C8 and C10 reveal critical residues (green spheres), shown on the ZIKV E ectodomain (PDB # 5IRE; (9).

To further test the native antigenicity of ZIKV RVPs, we performed an ELISA with RVPs using various conformational MAbs. As detection MAbs, we used two EDE MAbs (C8 and C10) and five neutralizing anti-ZIKV MAbs (ZIKV-117, ZIKV-195, ZIKV-116 (29), A9E (28), and LM-081). Each of these seven MAbs represents a distinct conformational region on the native structure of ZIKV prM/E (Fig. 1D). Two negative control MAbs were also tested that do not bind or neutralize ZIKV (CHIKV-specific CKV063 and DENV1-specific 1F4). The capture MAb was a mouse anti-fusion loop MAb (4G2). As expected, the anti-ZIKV MAbs showed strong levels of binding, while negative control MAbs did not bind ZIKV RVPs (**Fig. 2A**). The binding by the EDE MAbs C8 and C10 is particularly suggestive of ZIKV RVPs having native antigenicity, since these MAbs are known to bind complex, quaternary, epitopes on the flavivirus surface (39). To better define the epitope residues of LM-081, C8, and C10, we epitope mapped these MAbs using our Shotgun Mutagenesis Epitope Mapping platform (**Figs 2C** and **2D**, and **Table S2**). These results confirm that our ZIKV RVPs are antigenically indistinguishable from live ZIKV, as expected since ZIKV RVPs are made with the identical CprME protein from wild type ZIKV.

### ZIKV RVPs are stable and infectious after incubation at elevated temperature, freeze-thaw, or prolonged cryopreservation

The ability to detect ZIKV infectivity and neutralization can be limited by stability of the virus under commonly used experimental conditions, including incubation of up to 2 h with neutralizing antibodies or serum. To determine the stability of ZIKV RVP preparations during typical infection conditions, we incubated ZIKV RVPs at 4, 25, or 37°C for up to 72 hours, and then infected BHK-DC-SIGN cells. Our results show that RVPs remain infectious under typical incubation temperatures (4–37°C) and time periods (e.g., 1–2 h) used in neutralization assays (mean infectivity at 2 h of 96% at 4°C, 87% at 25°C, and 69% at 37°C) (**Fig. 3A**).

**Figure 3.**
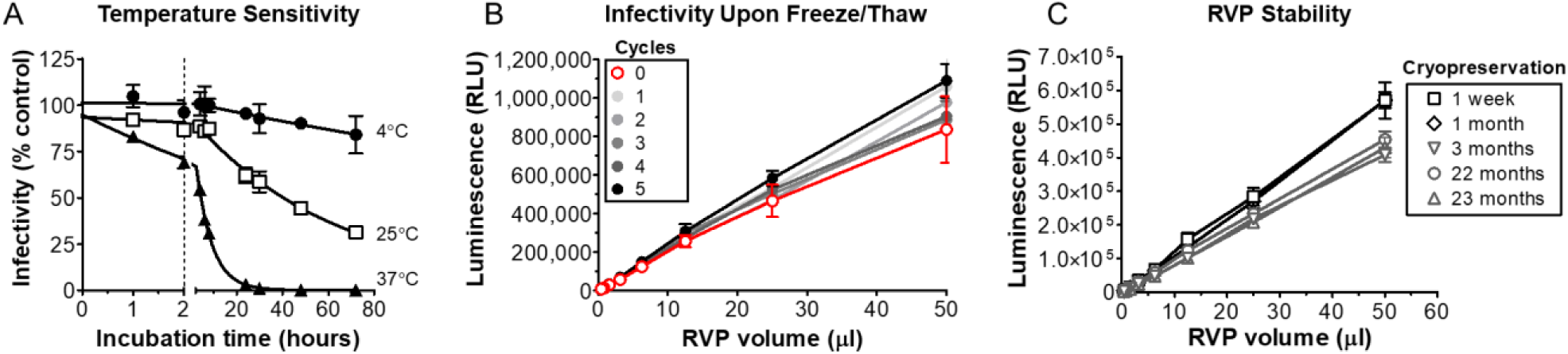
ZIKV RVPs are infectious and stable under commonly used conditions. **(A)** Cryopreserved ZIKV RVP aliquots were thawed, mixed, incubated at 4, 25, or 37°C, and used to infect BHK-DC-SIGN cells. Infectivity was compared to that of freshly thawed ZIKV RVPs. Each data point is the mean of 3 replicate wells. **(B)** Infectivity of ZIKV RVPs sampled immediately after production or after multiple freeze/thaw cycles. Each data point represents the mean of at least 4 replicate wells. RVP infectivity did not diminish upon freeze/thaw when tested from 0 to 5 cycles. **(C)** Five different lots of ZIKV RVPs (produced at various times over 23 months) were tested for infectivity using BHK-DC-SIGN target cells. RVPs were added to 96-well black plates and subjected to serial, 2-fold dilutions before adding target cells. Each data point is the mean of 8 replicate wells. All error bars represent SD.

ZIKV RVPs were also tested for their ability to infect cells immediately after production or after multiple freeze-thaw cycles. The results showed that ZIKV RVPs are robust, maintaining their ability to infect cells even after 5 rounds of refreezing and rethawing (**Fig. 3B**). Our results are consistent with the stability of ZIKV being high compared to DENV under equivalent conditions (42, 43).

A single lot of a critical reagent is often used for large-scale experiments that occur over an extended period of time. Thus the ability to store a single lot of ZIKV RVPs is important. To assess the stability of cryopreserved ZIKV RVPs, we assayed the infectivity of 5 lots of ZIKV RVPs stored at −80°C from 1 week to 23 months. Although some decrease in infectivity was observed after 1 month of storage, RVPs stored for 23 months remained stable with robust levels of infectivity (**Fig. 3C**). These results demonstrate that ZIKV RVPs can be a reliable reagent even for experimentation that occurs over a prolonged time period.

### ZIKV RVPs can be used to derive reproducible neutralization titers

To test the reproducibility of neutralization measurements obtained using ZIKV RVPs, we performed replicate neutralization assays using the ZIKV-neutralizing MAb C8 and four dilutions of a single lot of ZIKV RVPs. RVPs were reproducibly neutralized by C8 (**Fig. 4A**), and the 50% neutralization titer (NT_50_) values were not statistically different (**Fig. 4B**, one-way ANOVA, *p* = 0.68). These data show that ZIKV RVP preparations obey the laws of mass action (44), do not contain interfering excess antigen, and, thus, can be used to determine accurate neutralization titers. In addition, we observed similar NT_50_ values for independent neutralization experiments performed on 5 different days (**Fig. 4C**) and across independently derived lots of RVPs (**Fig. 4D**), highlighting the reproducibility and reliability of RVPs for neutralization assays.

**Figure 4.**
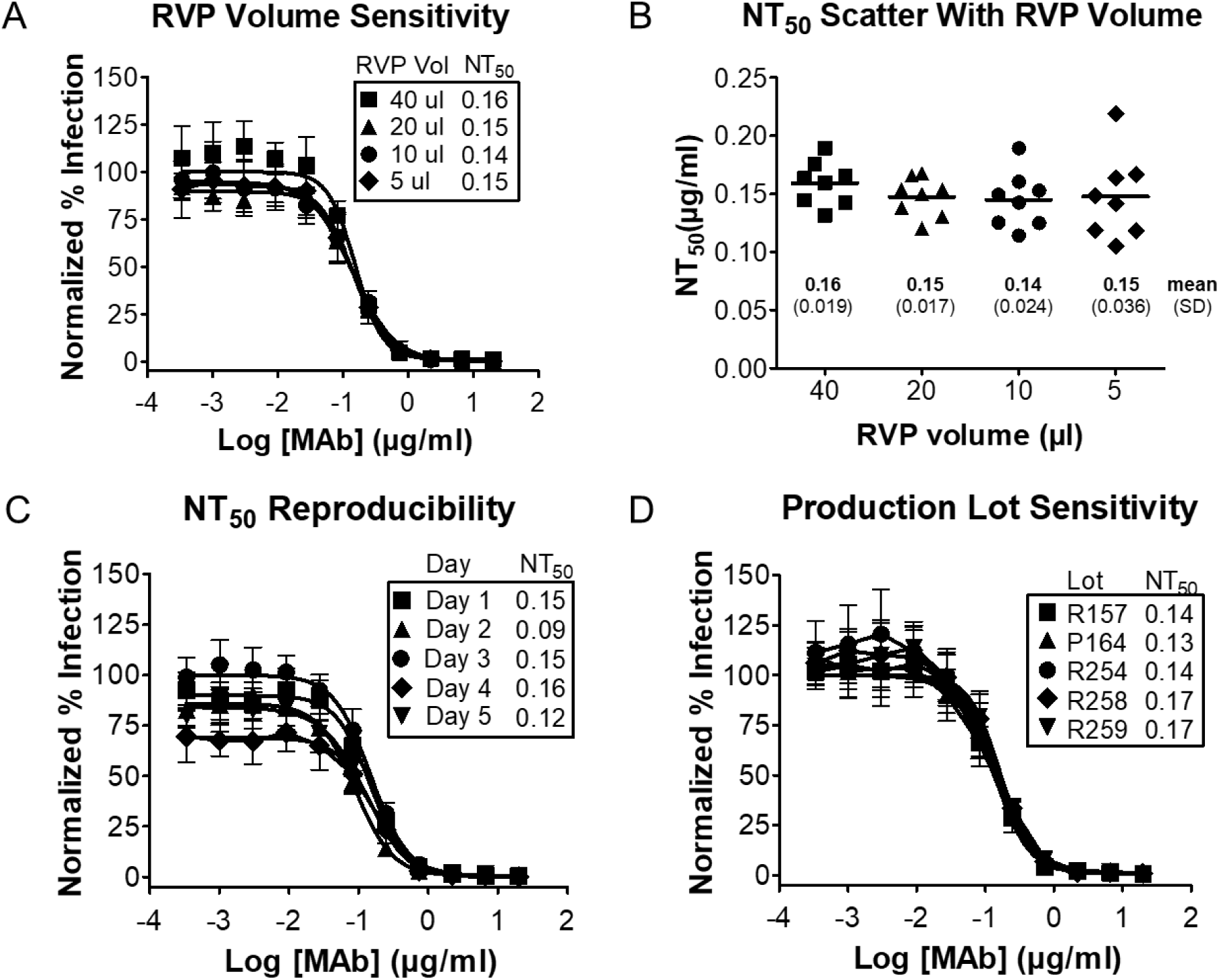
ZIKV RVPs can be used to derive reproducible antibody neutralization titers. **(A)** MAb C8 neutralization of RVPs was performed using various volumes (5 - 40 uL/well) of ZIKV RVPs from a single lot (#258A). RVPs were combined with serial 3-fold dilutions of C8 and incubated for 1 h at 37°C prior to infecting BHK-DC-SIGN cells. **(B)** NT_50_ values from A are plotted for each of the eight replicates run for each RVP volume. The mean NT_50_ value for each volume is shown by a horizontal line, and the mean NT_50_ value (μg/mL) and SD for each volume is as shown. **(C)** C8 neutralization of RVPs was performed on 5 different days, using one lot of ZIKV RVPs at 20 uL/well. **(D)** ZIKV neutralization by C8 was performed with 5 different lots of ZIKV RVPs, produced at various times over ∼2 years. Each lot was used at 20 uL/well. Data points for curves in **A-D** are means of 8 replicates, error bars are SD.

To be useful as a detection reagent for large-scale clinical ZIKV studies, ZIKV RVPs must be able to identify the activity of neutralizing antibodies in human serum. To test this ability, we used ZIKV RVPs in neutralization assays with serum from a naturally infected patient (Boca Biolistics). Similar to our MAb neutralization results, ZIKV RVPs were completely neutralized by the ZIKV-positive serum (**Fig. 5**). As expected, control (naïve) serum failed to neutralize any ZIKV RVPs.

**Figure 5.**
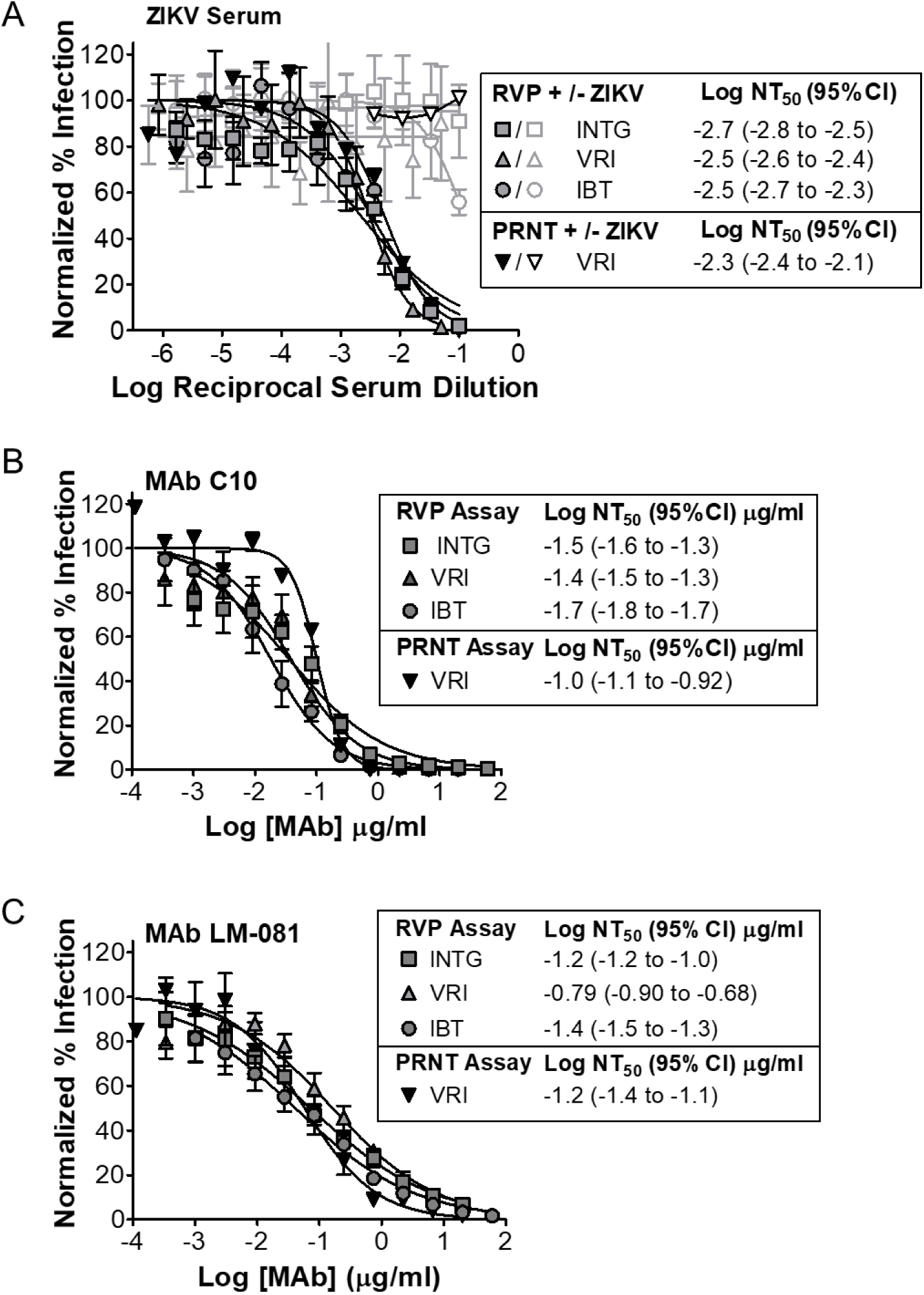
ZIKV RVPs are reproducibly neutralized in different laboratories by human serum from a ZIKV positive patient and by MAbs. **(A)** ZIKV RVP neutralization experiments using serum from a ZIKV infected individual (closed symbols) were conducted in three different laboratories: Integral Molecular (INTG), Vitalant Research Institute (VRI) and Integrated Biotherapeutics (IBT). Serum from an uninfected individual was used as a negative control (open symbols). Neutralization experiments were performed in different laboratories using MAbs **(B)** C10 and **(C)** LM-081. Data points for RVP neutralization assays represent the mean and standard deviation for 5-8 replicate data points. For PRNT assays, data points represent the mean and range of 2 replicates.

To further assess the reproducibility of neutralization data derived using ZIKV RVPs, experiments were independently performed at Integral Molecular alongside two other laboratories (Vitalant Research Institute [VRI] and IBT). Neutralization experiments tested both patient serum (Boca Biolistics) and MAbs (C10 and LM-081). The three laboratories independently obtained similar NT_50_ values, and in most cases these values had overlapping confidence intervals (**Fig 5**). PRNT NT_50_ values were comparable to those obtained using RVPs. In all cases, RVP and PRNT values were within 3-fold of the reference NT_50_ values obtained at Integral Molecular with ZIKV RVPs.

### Antibody-dependent enhancement

Antibodies that bind weakly or that do not neutralize DENV can increase pathogenesis in humans and animal models, in a process termed antibody-dependent enhancement (ADE). Sera and antibodies derived from DENV infection or from DENV vaccines are able to enhance ZIKV infection *in vitro* and *in vivo* (45, 46), although the impact of this enhancement on ZIKV pathogenesis is not clear. To demonstrate that our ZIKV RVPs can be used to detect ADE, we mixed serial dilutions of several ZIKV-reactive MAbs with ZIKV RVPs and added K562 cells, which naturally express the Fc-gamma receptor that provides an entry pathway for DENV or ZIKV virions bound by antibody. After 72 h, cells were tested for ZIKV infection (luciferase expression). All MAbs tested showed enhancement of infectivity, indicated by an increase in infection relative to the no-antibody control (**Fig. 6**), demonstrating the ability of our ZIKV RVP system to be used for the *in vitro* measurement of ADE.

**Figure 6.**
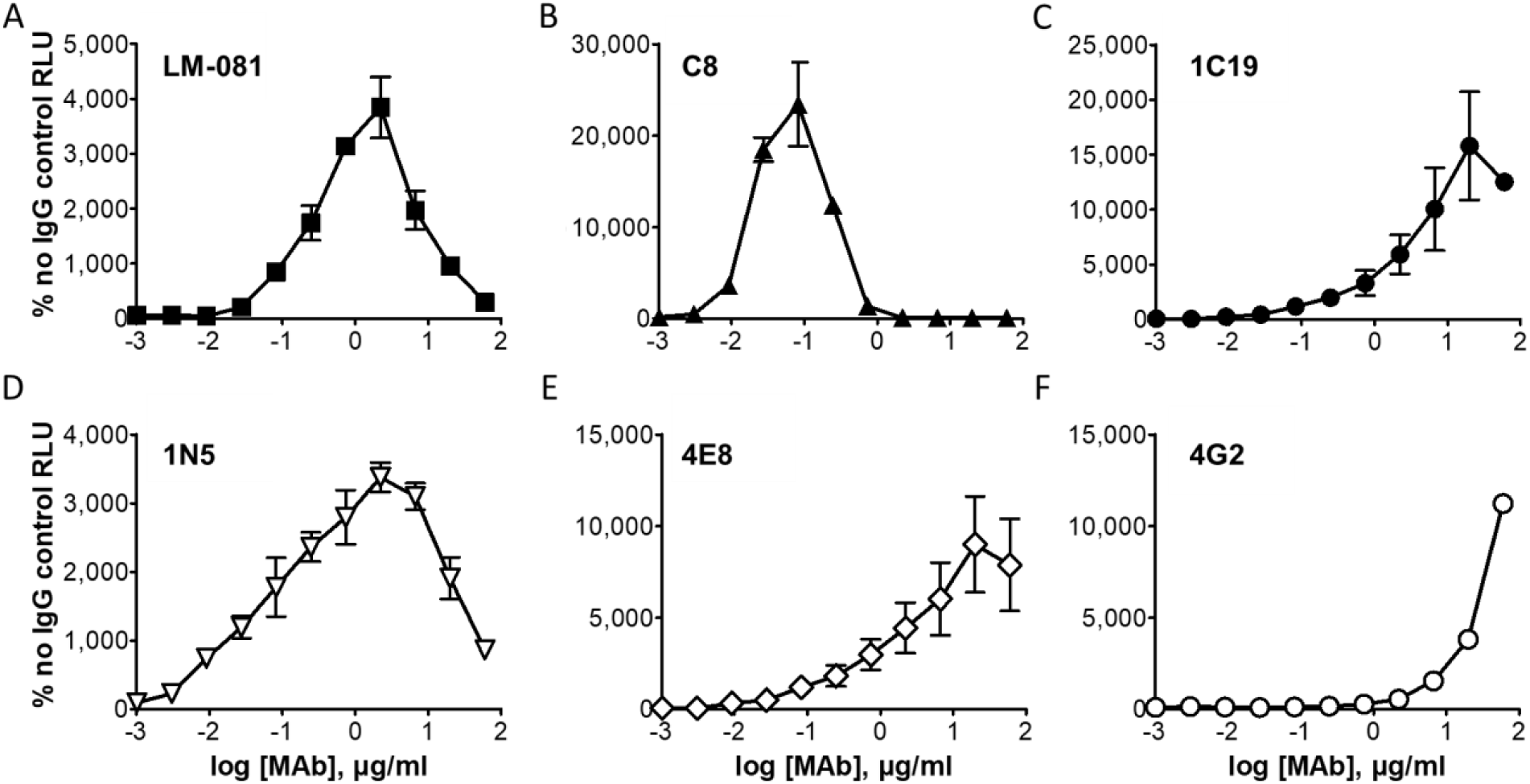
Antibody-dependent enhancement (ADE) of ZIKV infection. ZIKV RVPs were mixed and incubated for 1 h at 37°C with serial dilutions of ZIKV-reactive neutralizing MAbs LM081, C8, 1C19 (filled symbols) or non-neutralizing MABs 1N5, 4E8, 4G2 (unfilled symbols). K562 cells were then added to each well and incubated for 72 h at 37°C. Culture medium was removed, cells were lysed, and plates were analyzed for luciferase activity. Each data point is the average of 2 wells and is plotted as a percentage of the no antibody control. Error bars indicate range.

### Production of ZIKV RVPs

We have produced numerous lots of ZIKV RVPs for both small-scale (milliliter) and large-scale (liter) applications. We have not observed any significant differences in ZIKV RVP characteristics depending on production volume. Endotoxin, total aerobic microbial count, and total yeast and mold counts have all been below the limits of detection when tested (**Table S1)**.

To confirm that ZIKV RVPs do not replicate in cells, plaque formation was tested using two different cell types (Vero and BHK DC-SIGN) infected with ZIKV RVPs (1 ml RVPs/2×10(6) cells in a 6-well). No plaques were visible after cell growth with ZIKV RVPs for 4 days at 37°C. To further detect any viral replication, cell monolayers were fixed and stained for E protein using MAb 4G2 and a fluorescently labeled secondary antibody. No plaques were visible in cells grown with ZIKV RVPs, whereas plaques were visible in a live virus control well (**Fig. S2**). These results confirm that ZIKV RVPs are replication-incompetent under normal conditions of flavivirus infection and growth.

### Generation of ZIKV RVPs with distinct maturation states

Maturation of flaviviruses occurs by the cleavage of the prM protein to M (9), but this is generally incomplete, with virus maturity (whether live virus or RVP) (Cherrier 2009; ref for ZIKV) being influenced by the producing cell type (Dejninarattisai et al., 2015). In addition, patient-derived DENV showed higher levels of maturity compared with the same virus following passage in cells (47, 48). The maturation state of flaviviruses is thought to be important for infectivity and for neutralization by sera in clinical studies. To investigate the ZIKV RVP maturation state, and to demonstrate the ability to modify maturation if desired, we produced different lots of ZIKV RVPs either under standard conditions or in the presence of over-expressed (transfected) furin to promote RVP maturation by cleaving prM (9). Western blot analysis confirmed the near-complete cleavage of prM to M in RVPs produced in the presence of transfected furin, while prM was only partially cleaved in RVPs from standard production conditions (**Fig. 7A**), indicating partial RVP maturity. Both fully mature and partially mature ZIKV RVPs demonstrated robust infection (**Fig. 7B**), although infectivity of the more mature ZIKV RVPs was slightly lower. This decrease in infectivity was likely due to a slight reduction in particle production, as suggested by the signal for E protein by Western analysis. The neutralization of ZIKV RVPs with various MAbs was not altered by ZIKV maturation (**Fig. 7C**), suggesting that in RVPs produced under standard conditions the level of remaining prM did not affect the ability to be neutralized, even by EDE MAbs whose binding site overlaps with that of prM. Our results demonstrate the creation of both mature and partially mature ZIKV RVPs, and both are infectious and antigenically identical to wild type with respect to anti-E MAbs.

**Figure 7.**
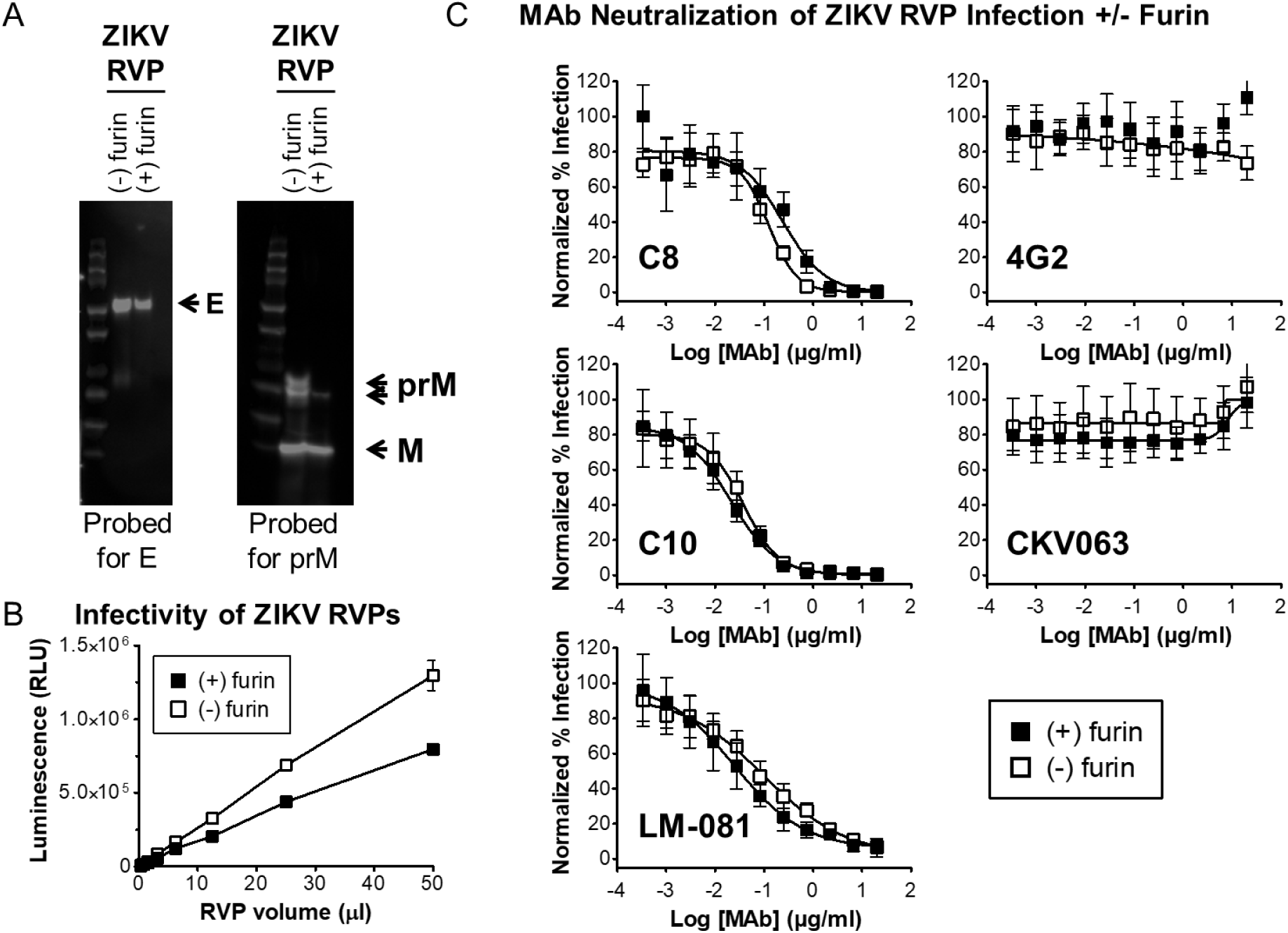
Control of ZIKV RVP maturation state. **(A)** Western blot showing maturation state of ZIKV RVPs produced in the absence or presence of transfected human furin. **(B)** Infectivity of ZIKV RVPs produced under normal conditions or in the presence of transfected furin. Each data point is the mean of 2 wells, error bars represent the range. **(C)** Neutralization of infectivity by ZIKV RVPs produced under normal conditions in the absence or presence of transfected furin. ZIKV RVPs were incubated with serial dilutions of MAbs C8, C10, LM-081, 4G2, and CKV063. Each data point represents the average of 8 wells, error bars are SD.

## DISCUSSION

We previously produced pseudo-infectious DENV RVPs that were designed to be antigenically identical to each of the four DENV serotypes. We adapted this system to ZIKV, developing ZIKV RVPs as a means for rapidly determining the neutralizing activity of ZIKV MAbs and sera. Our ZIKV RVPs have already been used to determine neutralizing antibody titers in immunized animals during preclinical studies of two different ZIKV vaccines (23, 24). Another study used our ZIKV and DENV RVPs for testing of longitudinal human serum samples following positive results by RT-PCR and ELISA, reporting the first cases of ZIKV-infected U.S. blood donors outside states with areas of active transmission (25). Other groups have used similar ZIKV RVPs in other infection and neutralization studies (42, 49). The use of RVPs as critical reagents in human clinical trials for vaccines and therapeutics requires their further validation using stability and reproducibility metrics for large-scale applications. Here, we describe the results of validation studies, demonstrating that our ZIKV RVPs display antigenic integrity, are stable under typical conditions used in infectivity assays, and can be reproducibly produced and used in neutralization assays that test MAbs and serum.

Our ZIKV RVPs were highly robust and thermostable, retaining their native antigenicity and infectivity after cryopreservation up to ∼2 years. The ZIKV RVPs also retained their infectivity after 5 freeze-thaw cycles. Although prolonged incubation (>4 h) at 37°C did decrease infectivity below 50%, ZIKV RVPs otherwise showed robust thermostability under typical conditions used in neutralization assays (e.g., 37°C for 1 hour). ZIKV has been reported to display significantly increased thermostability relative to DENV (42, 43), and our results are consistent with this enhanced stability. Our results suggest that ZIKV RVPs are antigenically equivalent to live ZIKV, which is expected since ZIKV RVPs are made with the identical CprME protein as live ZIKV. However, it is noted that ZIKV RVPs are made in defined cell types (typically BHK cells), and there is evidence suggesting that some flavivirus properties, particularly maturity, can be influenced by the producing cell type (whether live virus or RVP) (47).

To diagnose flavivirus infection, the U.S. Centers for Disease Control and Prevention (CDC) recommends a combination of molecular testing for genetic material (e.g., PCR) and confirmatory serologic testing, such as by PRNT (50). Although the PRNT assay has been the default test for measuring flavivirus neutralization for 50 years (15), the test has many disadvantages. For example, PRNT requires the use of live virus, which is associated with safety concerns. Certain viral strains may lead to false-negatives in the PRNT assay due to an inability of a strain to form plaques (16). Moreover, PRNT is a labor-intensive, highly variable, and technically complex assay that is not readily adaptable to high-throughput analysis of large numbers of clinical samples. For these reasons, in February of 2016, the U.S. FDA authorized the emergency use of molecular- and serological-based assays for ZIKV (51). To address the limitations of PRNT, companies have developed alternative assays for detecting ZIKV infection and neutralization, based on microneutralization, flow cytometry, and ELISA (52-54). These formats have several advantages over PRNT, including single-cell detection, a readout independent of plaque formation, and relatively rapid detection (2–5 days post-infection). However, many of these assays still rely on live infectious virus and cannot always be readily adapted to large-scale, high-throughput detection of neutralizing antibodies. Nevertheless, it should be noted that plaque formation assays can be used to measure multiple aspects of the viral life-cycle, including entry, viral egress, and viral spread between cells, which may be desirable measurements for some viruses and mechanisms of viral inhibition (55, 56).

RVPs have advantages and applications beyond the antibody neutralization assays tested here. RVPs can also be used to test or screen for small molecule inhibitors that have potential as drugs. Drugs that affect prM/E production and cleavage, cellular uptake, or attachment to cells could be identified and tested using RVPs. The use of RVPs in combination with functional genomic platforms can be used to identify new attachment factors and receptors, or changes within a cell that affects viral entry. Moreover, RVPs enable rapid studies of prM/E structure-function or the testing of specific mutations or strains that have desirable properties, including variants that are not capable of replication and therefore could not be studied using live virus (57). Purified RVPs can also be used in antibody characterization studies, including epitope mapping, ELISA, and biosensor kinetic studies (58).

Most of the MAb epitope maps reported here have been published elsewhere previously. However, the epitopes of the EDE MAbs C8 and C10 are of particular note because they are highly conformationally complex and require dimerized E molecules to bind. The epitope of C8 on ZIKV prM/E has been reported previously (39) and agrees with our epitope map derived by shotgun mutagenesis alanine scanning, which identifies the residues with the greatest energetic contribution to binding. The epitope of C10 on ZIKV has not been published previously. The commercial ZIKV-specific MAb LM-081 has not been published previously, but neutralizes with similar potency to C8 and C10 (NT_50_ of 72 ng/ml vs. 123 and 36 ng/ml, respectively). LM-081 appears to derive its specificity by binding E protein Domain III residue I317, which is highly specific to ZIKV (the equivalent position in all 4 DENV serotypes is a glutamic acid).

Our results demonstrate that ZIKV RVPs provide a safe, fast, quantitative, and highly reproducible platform for measuring ZIKV infectivity and identifying neutralizing and enhancing antibodies against ZIKV. The ability of ZIKV RVPs to accurately detect antibodies and neutralizing activity in patient serum highlights their utility for clinical applications. ZIKV RVPs may serve as an attractive alternative to existing methodologies, with particular application to large-scale studies such as ZIKV vaccine trials, epidemiologic surveillance, and high-throughput drug screening.

## ACKNOWLEDGEMENTS

This work was supported by NIH contract HHSN272201400058C to BJD. GS was supported by NIH grant R01 AI119056. We thank Soma Banik, Duncan Huston-Paterson, and Andrey Efimov for valuable technical assistance. We also thank Hetal Patel, Hansi Dean, Steph Sonnberg, and Swati Mukherjee for valuable scientific advice.

## TABLES

**Table S1.** Quality control tests and results for luciferase ZIKV RVPs.

| Test | Result |
| --- | --- |
| Bioburden (USP61) | < 1 cfu/mL TYMC<sup>a</sup> |
| Bioburden (USP61) | < 2 cfu/mL TAMC<sup>b</sup> |
| Endotoxin | < 1 EU/mL<sup>c</sup> |
| *Mycoplasma* (PCR) | Negative<sup>d</sup> |
| *Mycoplasma* (ELISA) | Negative<sup>e</sup> |

<sup>a</sup> TYMC: total yeast and mold count (limit of detection : 1 cfu/mL)
<sup>b</sup> TAMC: total aerobic microbial count (limit of detection: 2 cfu/mL)
<sup>c</sup> GenScript ToxinSensor Gel Clot Endotoxin Assay System (limit of detection: 1 EU/mL)
<sup>d</sup> Sigma LookOut PCR kit, result indicated by absence/presence of specific PCR product
<sup>e</sup> Lonza MycoAlert kit, result interpreted according to manufacturer's protocol

**Table S2.** Epitope mapping data for MAbs LM-081, C8 and C10. MAb reactivities for each alanine scan mutant are expressed as a percentage of reactivity with wild-type ZIKV prM/E, with ranges (half of the maximum minus minimum values) in parentheses. Values for critical residues are shaded in gray. Values shown are the average of at least two replicate experiments. Data for anti-ZIKV MAbs A9E, ZIKV-116, and ZIKV-117 are also shown as comparative controls.

| Residue Mutation | Test MAbs |  |  | Control MAbs |  |  |
| --- | --- | --- | --- | --- | --- | --- |
|  | C10 | C8 | LM-081 | A9E | ZIKV-116 | ZIKV-117 |
| R2A | 5 (2) | 29 (5) | 43 (6) | 84 (11) | 116 (23) | 70 (2) |
| R73A | 28 (5) | 56 (3) | 92 (1) | 106 (28) | 121 (13) | 53 (9) |
| G78A | 72 (8) | 9 (2) | 41 (12) | 81 (5) | 97 (10) | 18 (3) |
| R99A | -1 (2) | 0 (1) | 53 (5) | 61 (6) | 108 (14) | 53 (9) |
| N103A | -1 (1) | 0 (1) | 25 (9) | 59 (1) | 96 (20) | 35 (3) |
| G104A | 19 (2) | 8 (1) | 41 (4) | 55 (6) | 84 (4) | 35 (2) |
| G106A | 160 (20) | 72 (2) | 22 (5) | 112 (14) | 120 (16) | 140 (28) |
| G109A | 7 (-) | -1 (1) | 74 (6) | 73 (5) | 104 (7) | 45 (4) |
| L113A | 105 (5) | 2 (3) | 98 (17) | 92 (10) | 90 (7) | 61 (11) |
| V153A | 24 (9) | 73 (3) | 117 (30) | 116 (3) | 133 (15) | 117 (14) |
| R252A | 193 (43) | 12 (2) | 101 (62) | 112 (8) | 111 (11) | 156 (15) |
| K316A | -2 (3) | 1 (3) | 42 (12) | 75 (1) | 75 (0) | 103 (15) |
| I317A | 73 (10) | 111 (26) | 13 (5) | 85 (12) | 109 (7) | 122 (5) |
| K373A | 42 (7) | 24 (4) | 80 (19) | 58 (11) | 47 (0) | 74 (11) |

## FIGURES AND LEGENDS

**Figure S1.**
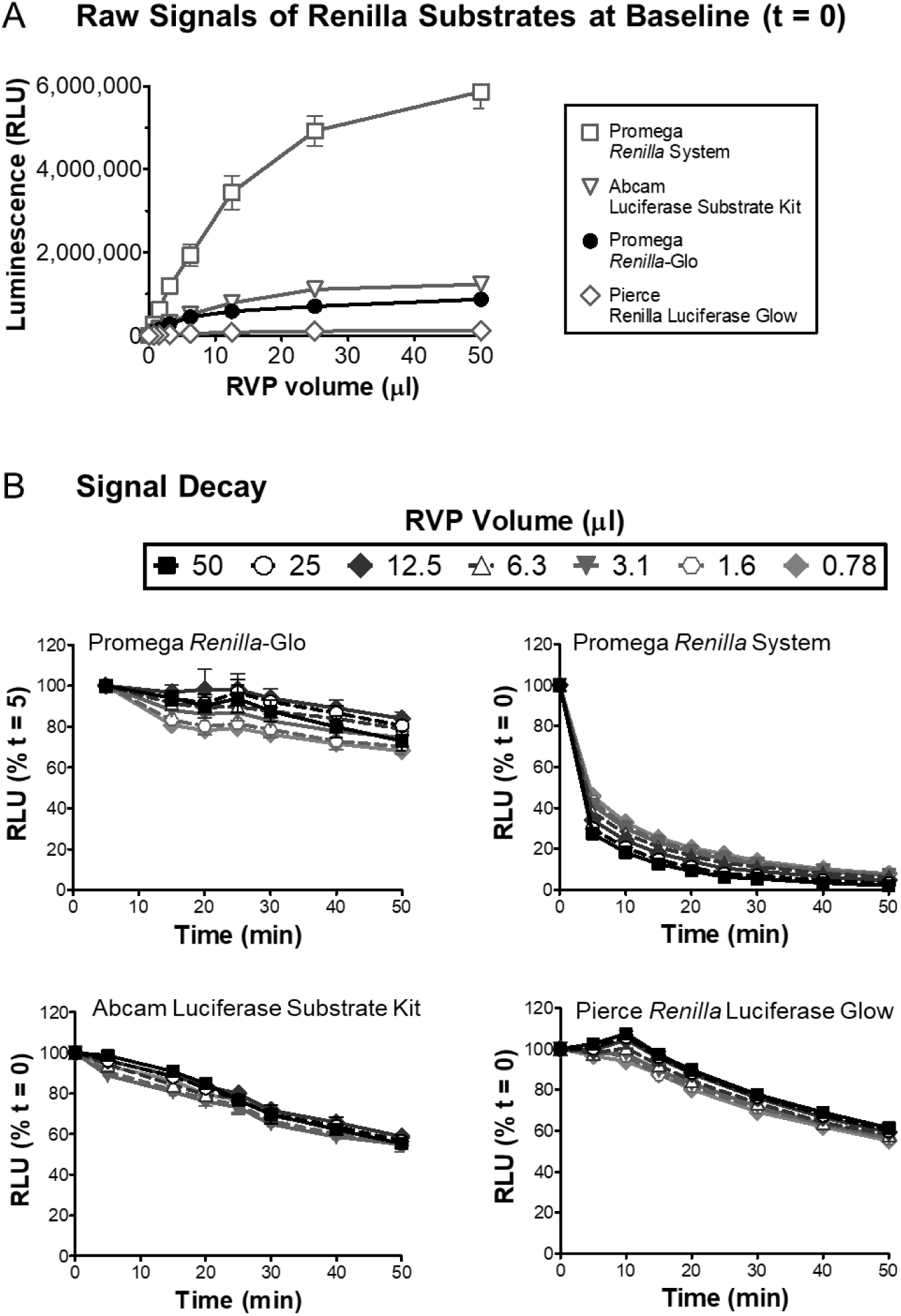
Use of RVPs with different *Renilla* substrates. BHK-DC-SIGN cells infected with various volumes of luciferase RVPs were lysed and mixed with different luciferase substrates according to the manufacturers’ instructions. **(A)** Luminescence from individual samples (in relative luminescence units, RLU; n = 3, error bars are SD) was detected on an Envision plate reader after adding substrate. **(B)** Luminescence from individual samples was detected over time and plotted as a percentage of the signal at time 0 min (n = 3, error bars are SD). Luciferase assay reagents tested included the Promega *Renilla*-Glo Luciferase Assay System (E2710) (recommended), Promega *Renilla* Luciferase Assay System (E2810), Abcam Luciferase Reporter Assay Substrate Kit (ab228546), and Pierce *Renilla* Luciferase Glow Assay Kit (ThermoFisher 1616). For Promega *Renilla*-Glo (top left panel), luminescence was variable immediately after addition of substrate but stabilized after 5 min, so data was normalized to the 5 min time point rather than to 0 min.

**Figure S2.**
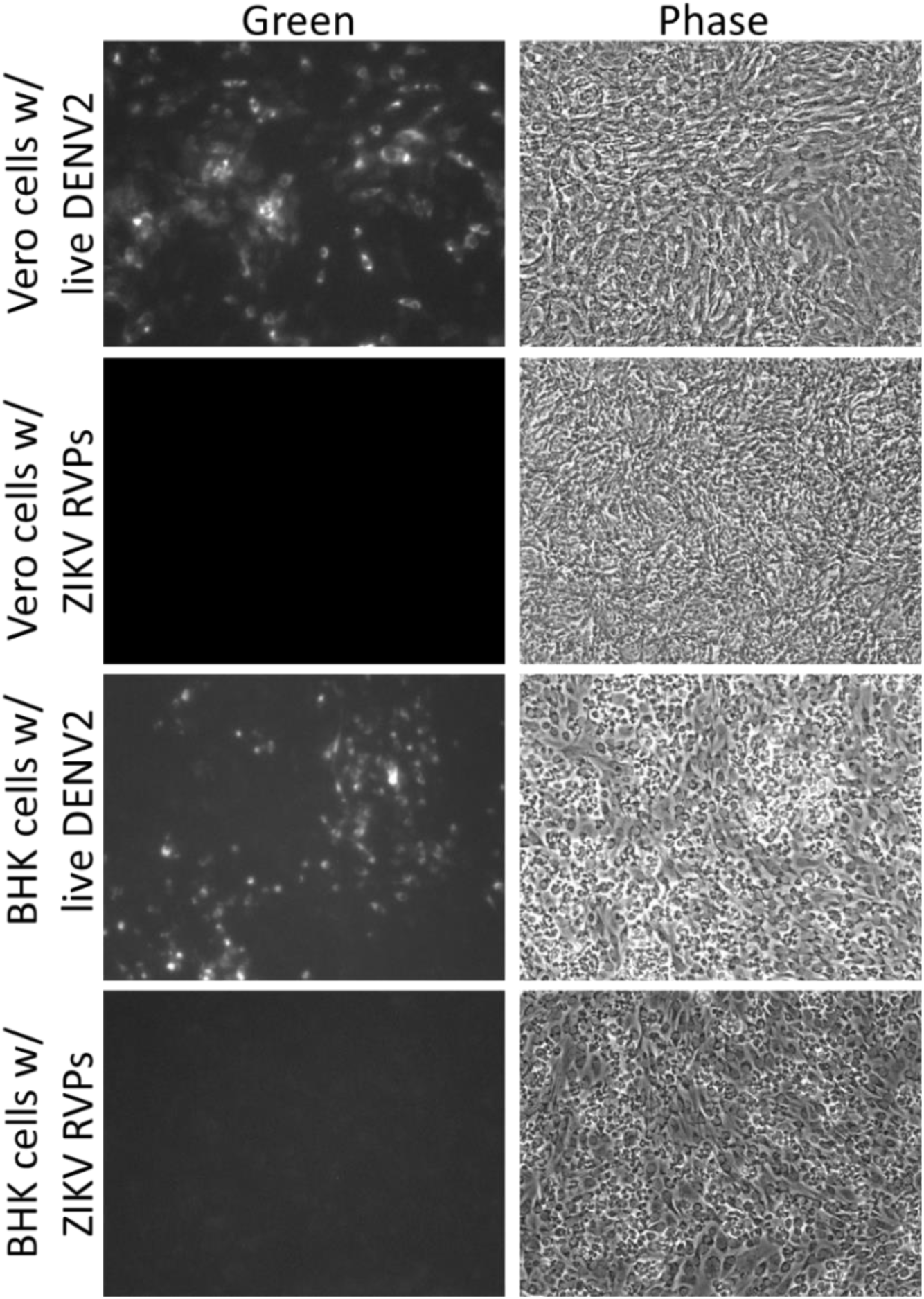
Plaque formation assays. Vero or BHK DC-SIGN cells were seeded in 6-well tissue culture plates and grown for 1 d. Medium was replaced with ZIKV RVPs (1 mL/well) or live DENV2 virus in BHK Infection Medium. After 1 h at 37°C, 1 mL of medium was added and cells were grown for 4 d (37°C, 5% CO_2_). Cell monolayers were fixed, blocked, and stained for flavivirus E protein using MAb 4G2 (2 ug/mL) and goat anti-mouse Alexa Fluor 488. Fluorescence (**Green**) and phase-contrast microscopy (**Phase**) results are shown.

